# Discovering single nucleotide variants and indels from bulk and single-cell ATAC-seq

**DOI:** 10.1101/2021.02.26.433126

**Authors:** Arya R. Massarat, Arko Sen, Jeff Jaureguy, Sélène T. Tyndale, Yi Fu, Galina Erikson, Graham McVicker

## Abstract

Genetic variants and de novo mutations in regulatory regions of the genome are typically discovered by whole-genome sequencing (WGS), however WGS is expensive and most WGS reads come from non-regulatory regions. The Assay for Transposase-Accessible Chromatin (ATAC-seq) generates reads from regulatory sequences and could potentially be used as a low-cost ‘capture’ method for regulatory variant discovery, but its use for this purpose has not been systematically evaluated. Here we apply seven variant callers to bulk and single-cell ATAC-seq data and evaluate their ability to identify single nucleotide variants (SNVs) and insertions/deletions (indels). In addition, we develop an ensemble classifier, VarCA, which combines features from individual variant callers to predict variants. The Genome Analysis Toolkit (GATK) is the best-performing individual caller with precision/recall on a bulk ATAC test dataset of 0.92/0.97 for SNVs and 0.87/0.82 for indels. On bulk ATAC-seq reads, VarCA achieves superior performance with precision/recall of 0.99/0.95 for SNVs and 0.93/0.80 for indels. On single-cell ATAC-seq reads, VarCA attains precision/recall of 0.98/0.94 for SNVs and 0.82/0.82 for indels. In summary, ATAC-seq reads can be used to accurately discover non-coding regulatory variants in the absence of whole-genome sequencing data and our ensemble method, VarCA, has the best overall performance.

## INTRODUCTION

Rare genetic variants and somatic mutations in regulatory regions are important for many human traits and diseases [1], but typically require whole genome sequencing (WGS) to discover since they are not captured by single-nucleotide polymorphism (SNP) arrays and cannot be accurately imputed. WGS is expensive to perform on large panels of individuals, and the majority of sequencing reads come from non-coding and non-regulatory regions of the genome that are less likely to contain functional variants. While capture panels can be designed to target regulatory regions, the positions of active regulatory regions differ substantially across cell types, so very large or custom capture panels for each cell type would be required to survey all variants in regulatory regions.

One way to overcome these challenges is to identify genetic variants using data from the Assay for Transposase-Accessible Chromatin using sequencing (ATAC-seq) [2]. ATAC-seq generates sequence reads from regulatory regions of the genome and can potentially act like a low-cost capture method for important non-coding sequences. Several methods have been developed to genotype or call variants from chromatin immunoprecipitation followed by sequencing (ChIP-seq) data [3–5]. Previous research has evaluated the performance of single nucleotide variant detection methods applied to single-cell RNA-seq data [6]. However, the performance of single nucleotide variant (SNV) and insertion/deletion (indel) callers on bulk and single-cell ATAC-seq data has not been systematically evaluated. Furthermore, some features of ATAC-seq reads may diminish the performance of standard tools for variant detection. For example, ATAC-seq insert sizes are short and differ between nucleosome-free and nucleosome-spanning sequence fragments [2]. This is likely to cause problems for indel callers such as Manta [7], Pindel [8], and DELLY [9], which utilize the distribution of insert sizes from mapped reads. Furthermore, variant callers may fail to identify heterozygous variants when ATAC-seq reads originate primarily from one allele.

Here we compare the performance of several SNV and indel callers on bulk and single-cell ATAC-seq reads, using high quality genotypes from the Platinum Genomes project as a ground truth dataset [10]. We also develop an ensemble method, called VarCA for “variants in chromatin accessible regions,” which combines features from multiple callers. VarCA uses a random forest to predict indels and SNVs and achieves substantially better performance than any individual caller.

## MATERIAL AND METHODS

### VarCA method overview

VarCA is implemented as a Snakemake [11] pipeline with two components, which we refer to as the *prepare* and *classify* subworkflows (Fig 1). The *prepare* subworkflow runs multiple variant callers on aligned ATAC-seq reads and gathers the output of these callers together into a single dataset in variant call format (VCF). The *classify* subworkflow uses the output from the *prepare* subworkflow to predict variants within ATAC-seq peaks and outputs a new VCF file containing the predictions. In addition to the predicted variants, this VCF file contains recalibrated quality scores and the names of the variant callers that predicted each variant. By default, the two subworkflows are executed together in a master Snakefile, with the output of the first passed directly to the second, however, for more advanced usage, they can be executed separately. This allows the user to alter the output of the *prepare* subworkflow before it is used by the *classify* subworkflow, or to use the output of the *classify* subworkflow to evaluate the performance of the variant calling methods. The pipeline is highly configurable and allows the user to easily include additional variant callers.

**Figure 1.**
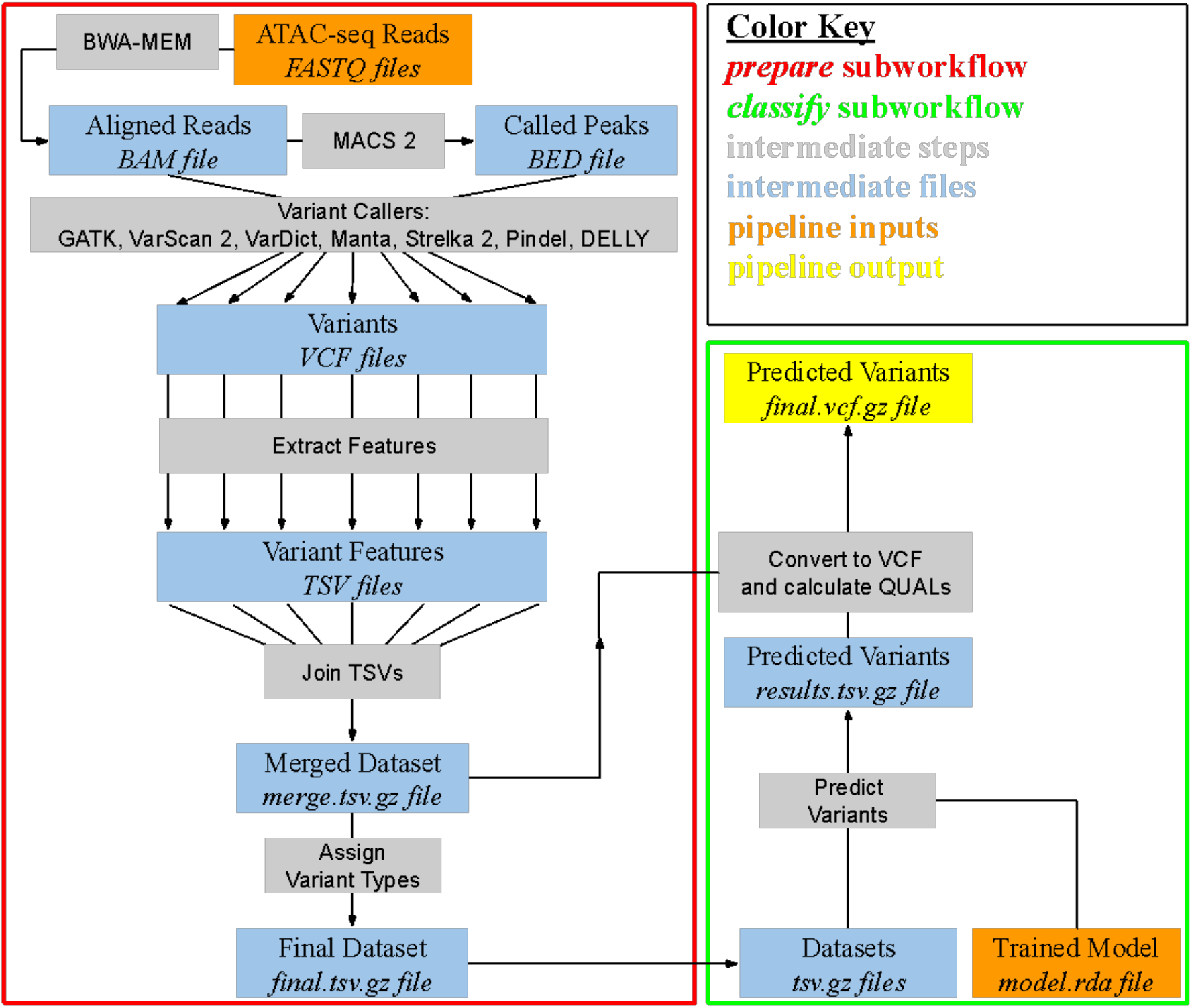
Schematic of the VarCA pipeline. The pipeline consists of the *prepare* (red) and *classify* (green) subworkflows. The *prepare* subworkflow runs a set of variant callers on the provided ATAC-seq data and extracts specific features from their VCF output. These are aggregated into a single dataset and prepared as input for the *classify* subworkflow, which uses a trained ensemble classifier (orange) to predict the existence of variants. The final output of the pipeline is a VCF file containing the variants predicted by VarCA (yellow).

### ATAC-seq data processing

The *prepare* subworkflow first aligns paired-end reads to a specified reference genome using BWA-MEM (version 0.7.15-r1140) [12] with default parameters. Reads are filtered with samtools (version=1.9) [13] and non-duplicated reads with high mapping quality (MAPQ ≥ 20) are retained. If the user desires, these steps of the pipeline can be skipped by providing pre-existing read alignments in BAM file format.

VarCA identifies ATAC-seq peaks using MACS2 (version 2.1.0.20150731) [14] with the following parameters’ --nomodel --extsize 200 --slocal 1000 --qvalue 0.05 -g hs’. This set of peak-calling arguments is more permissive and yields broader peaks than the ATAC-seq standard adopted by the ENCODE project. We use this peak definition because a more permissive peak definition is desirable for calling as many relevant variants as possible. The user may also provide their own peak definitions by providing a BED file, in which case the peak-calling step of the pipeline is skipped.

### Single cell ATAC-seq data processing

Single cell ATAC-seq reads from a mixture of human GM12878 and mouse A20 cells were obtained from the 10x Genomics website [15]. Reads were aligned to the hg19/mm10 reference genome and clustered according to the 10x Single Cell ATAC pipeline. Reads for each of the clusters were extracted to separate BAM files. Read pairs aligning to the mm10 reference genome were discarded with samtools (version=1.9) [13]. Alternative alignment locations mapping to the mm10 reference were also discarded using pysam [13]. Peak-calling for each cluster was performed identically to the bulk cell ATAC-seq. Only clusters with a vast majority of reads aligned to the human genome rather than the mouse genome were used by VarCA.

### Running indel and SNV callers

To call SNVs in the *prepare* subworkflow, VarCA runs the Genome Analysis Toolkit (GATK) [16], VarDict [17], and VarScan 2 [18]. To call indels, VarCA uses the same callers, and additionally uses Manta [7], Strelka2 [19], Pindel [8], and DELLY [9]. The command line arguments and software versions for these callers are summarized in Supplementary Table S1. Variants from each caller are left-aligned and normalized using bcftools [20]. When callers do not output values for a site, VarCA uses appropriate defaults specified in a separate configuration file. VarCA combines the output from the callers into a large feature table where each row represents a single genome position, and the columns (or features) are fields from the VCF output of each variant caller. As additional features, VarCA adds columns to the table with the depth of reads overlapping each position as well as Bayesian estimates of the proportion of reads with insertions and deletions (see Supplementary Note S1). Columns are also added to label the variants predicted by the callers as’ INS’, ‘DEL’,’ SNV’, or’.’ (for no variant) based on the reference and alternate alleles output by each caller (Supplementary Fig S1).

### Training/test data

To create a ground truth dataset of known variants in ATAC-seq peaks, we downloaded high-quality genotype calls for the GM12878 cell line from the Platinum Genomes project [10], and downloaded bulk ATAC-seq data from the same cell line [2]. We aligned the ATAC-seq reads to the genome using the *prepare* subworkflow described above and labeled sites within ATAC-seq peaks (with read depth greater than 10) as’ INS’, ‘DEL’,’ SNV’, or’.’ (for no variant). In total, the feature table for the GM12878 cell line consisted of 4,866,073 sites within ATAC-seq peaks with a read depth greater than 10 (Supplementary Fig S2). We partitioned the feature table into training and test sets by dividing it into odd- and even-numbered chromosomes, respectively.

### VarCA training

VarCA’s *classify* subworkflow uses a random forest (RF) to predict whether a genome site contains a variant. Specifically, VarCA utilizes the ranger fast RF implementation [21] from within the mlr package [22]. The RF has several hyperparameters that control the structure of the trained RF: *num.trees* (number of trees to grow), *mtry* (number of variables to possibly split at each node in each tree) and *min.node.size* (the minimum number of observations to retain in terminal nodes, which controls the depth of the trees). We set *num.trees* to 500 and estimated the other hyperparameters by testing 35 parameter combinations and performing 5-fold cross validation on the training dataset (Supplementary Figs S3 and S4). We selected the parameter values that yielded the highest F-beta score, although we note that the performance was relatively insensitive to the choice of these parameters. To compute F-beta, we used a beta value of 0.5 to give precision a higher weight than recall. We trained separate RFs to classify indels from non-indels and SNVs from non-SNVs. In total we used 39 features from 3 callers to predict SNVs and 65 features from 7 callers to predict indels (Supplementary Tables S2 and S3).

### VarCA output

VarCA’s final output is a set of predicted variants in the variant call format (VCF). The RF used by VarCA provides probabilities that a variant exists but does not provide the alternate allele. VarCA obtains the alternate allele from one of the individual callers that predicted the existence of the variant. Since the predicted alternate allele may occasionally differ between callers, VarCA takes the first alternate allele that is predicted by a caller, with the ordering of the callers specified in a configuration file. Currently, we prioritize callers based on their individual F-beta scores (Supplementary Tables S4 and S5). VarCA indicates the callers used to determine the alternate allele for each variant within the VCF output file.

Using the classification probabilities output by the random forest, we also calculate a recalibrated quality score for every variant in the VCF. This score is the phred-scaled probability that the variant was correctly classified (i.e. that an alternate allele exists at this position). We calculated the quality scores by binning sites in the test dataset by their RF classification probabilities and fitting a linear model between the phred-scaled RF classification probabilities and empirical precision of each bin (Supplementary Fig S5). VarCA uses this model to calculate the final quality scores for every variant in the VCF.

### ATAC-seq experiments

To test whether VarCA can detect known mutations using ATAC-seq data, we performed ATAC-seq experiments for the Jurkat, MOLT-4, CCRF-CEM, RPMI-8402 and K-562 cell lines using the Omni-ATAC-seq method [23], with minor modifications. In each experiment, 1×10^5^ cells were centrifuged at 1000 × g for 10 min at 4 °C. Following aspiration, a cell count of the supernatant was performed, the remaining cell number was calculated, and all further reagents in the protocol were titrated to this cell number. For every 5×10^4^ cells, nuclei were isolated in 50 μl cold ATAC-Resuspension Buffer (RSB) (10 mM Tris-HCl pH 7.4, 10 mM NaCl, 3 mM MgCl2) containing 0.1% NP40, 0.1% Tween-20, and 0.01% Digitonin, and pipet-mixed up-and-down at least 5 times. Nuclei isolation mix was incubated on ice for 3 exactly minutes, washed in 1 ml of cold ATAC-RSB containing 0.1% Tween-20 (but no NP40 or Digitonin) and centrifuged at 1000 × g for 10 min at 4 °C. Nuclear DNA was tagmented in 50 μl Transposition mix (25 μl 2 × TD buffer, 2.5 μl transposase (100nM final), 0.5 μl 1% digitonin, 0.5 μl 10% Tween-20, 16.5 μl PBS and 5 μl diH2O), and incubated in a thermomixer at 37 °C, 1000 × g for 30 min. Tagmented DNA was purified with Zymo DNA Clean and Concentrator-5 Kit (cat# D4014). Library amplification was performed using custom indexing Nextera primers from IDT in a 50 μl Kapa Hi Fi Hot Start PCR reaction (cat# KK2602). Following 3 initial cycles, 1 μl of PCR reaction was used in a quantitative PCR (Kapa qPCR Library Quantitation Kit cat# KK4824) to calculate the optimum number of final amplification cycles. Library amplification was followed by SPRI size selection with Kapa Pure Beads (cat# KK8002) to retain only fragments between 80-1,200bp. Library size was obtained on an Aglient Bio-Analyzer or TapeStation using a High Sensitivity DNA kit and factored into final Kapa qPCR results to calculate the final size-adjusted molarity of each library. Libraries were pooled and sequenced on an Illumina NextSeq500 in Paired-End 42 base pair configuration at the Salk Next Generation Sequencing Core. ATAC-seq data quality was assessed by computing the fraction of reads within peaks (FRiP) (Supplementary Table S6).

## RESULTS

### Assessing variant caller performance

To evaluate the performance of variant callers on ATAC-seq data, we considered all sites within ATAC-seq peaks with read depth greater than 10 and used Platinum Genomes variant calls to label classifications as false positives, true positives, false negatives, or true negatives. Since most variant callers only provide output for sites that they classify as variants, we used the variants output from each caller as predicted positives and the remaining sites as predicted negatives. Using this approach, we obtained point estimates of precision, recall and other performance metrics for each variant caller (Fig 2; Supplementary Tables S4 and S5).

**Figure 2.**
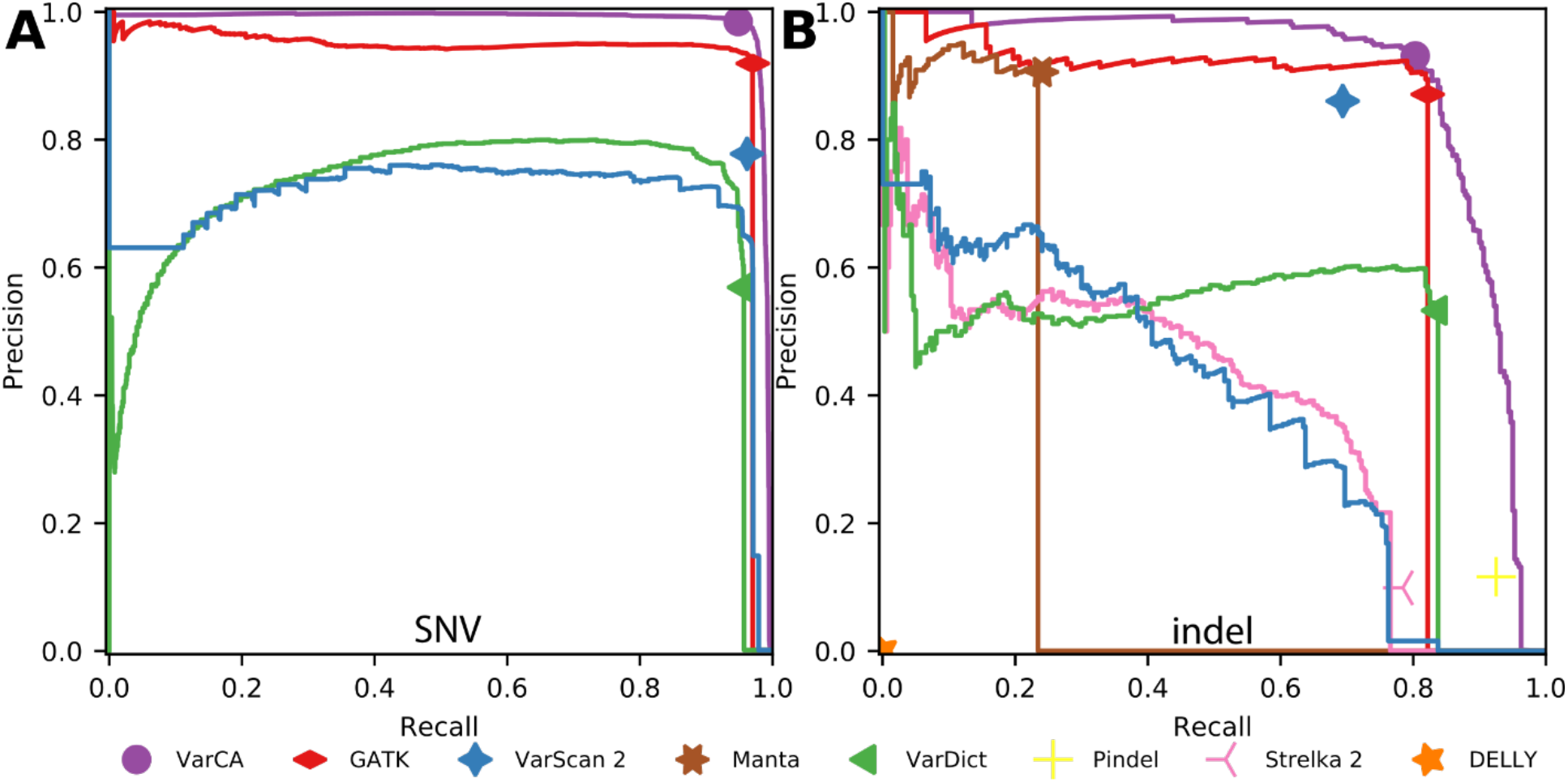
Performance of methods for detecting single nucleotide variants (SNVs) and indels on bulk ATAC-seq. (**A**) Precision recall (PR) curves for SNV detection (**B**) PR curves for indel detection. VarCA is our ensemble prediction method that combines information from individual variant callers. Overall precision and recall values for each caller, using all variant calls reported significant, are also indicated as single points. Callers that do not output a score that can be used for ranking only have a single point. The point for VarCA corresponds to a score of 0.50, which we use as a significance threshold. For each caller, sites that were not output/scored were assumed to be classified as non-variant and given a score lower than all output sites. The point for VarScan is substantially better than its precision recall curve because VarScan uses several metrics to decide which variants are significant and our VarScan ranking is based on a single metric (1-p-value).

We also generated precision-recall (PR) curves for the variant callers that provided scores that could be used to rank predictions. For example, for GATK we computed a PR curve using QD as the score (phred-scaled variant call confidence normalized by allele depth), for VarScan2 we used (1-p-value) as the score, and for VarDict, Manta, and Strelka, we used QUAL as the score (unnormalized phred-scaled variant call confidence). For each caller, we assigned unscored sites a value that was lower than all of the scored sites (the transition from scored to un-scored sites is visible as a sharp drop in precision in the PR plots). Some callers, such as Pindel and DELLY, do not output scores, so their performance is represented only by a single point on the PR plot. To assess the overall performance of each method we also computed the average precision over each curve, the area under the receiver operating characteristic curve (AUROC), and the F-beta score (using a beta value of 0.5) (Supplementary Tables S4 and S5).

Among the individual variant callers, GATK had the best performance for SNV discovery, as measured by F-beta, followed by VarScan2 and VarDict (Fig 2; Supplementary Tables S4 and S5). GATK also exhibited the best performance for indels followed by VarScan 2, Manta and VarDict. The remaining indel callers all had poor performance, likely due to their reliance on features, such as insert size, that have different characteristics in ATAC-seq reads.

VarCA, our ensemble classification method, had substantially better performance than the individual variant callers for both SNVs and indels, as measured by F-beta and had higher precision across all recall thresholds (Fig 2; Supplementary Tables S4 and S5). When run on SNVs for bulk cell ATAC-seq, VarCA had excellent performance with a precision of 0.99 and a recall of 0.95 (Supplementary Table S4) compared to a precision of 0.92 and recall of 0.97 for GATK. When run on indels for bulk cell ATAC-seq, VarCA had a precision of 0.93 and a recall of 0.80 compared to a precision of 0.87 and recall of 0.82 for GATK (Supplementary Table S5). In summary, by combining information from multiple variant callers, VarCA achieves substantially higher precision than GATK and all other variant callers, which can greatly reduce the number of false positive variant calls. For example, in our test dataset, VarCA reported 2983 true positive SNVs and only 46 false positive SNVs. In the same dataset GATK found 1.02 times more true positives than VarCA (3052) but at the cost of 4.8 times more false positives (268).

We next ran the individual variant callers and VarCA on single-cell ATAC-seq data, applying the methods separately to each cluster of cells. The overall performance of the methods on single-cell data was comparable to bulk data, and VarCA consistently had the highest precision (Fig 3; Supplementary Fig S6). For example, when applied to the reads from cells in “cluster 12”, VarCA had a precision of 0.98 and recall of 0.94 for SNVs and a precision of 0.82 and a recall of 0.82 for indels (Supplementary Table S6). This precision is higher compared to all other SNV and indel callers (Supplementary Table S7 and S8) and is consistent across the different cell clusters (Fig 3C).

**Figure 3.**
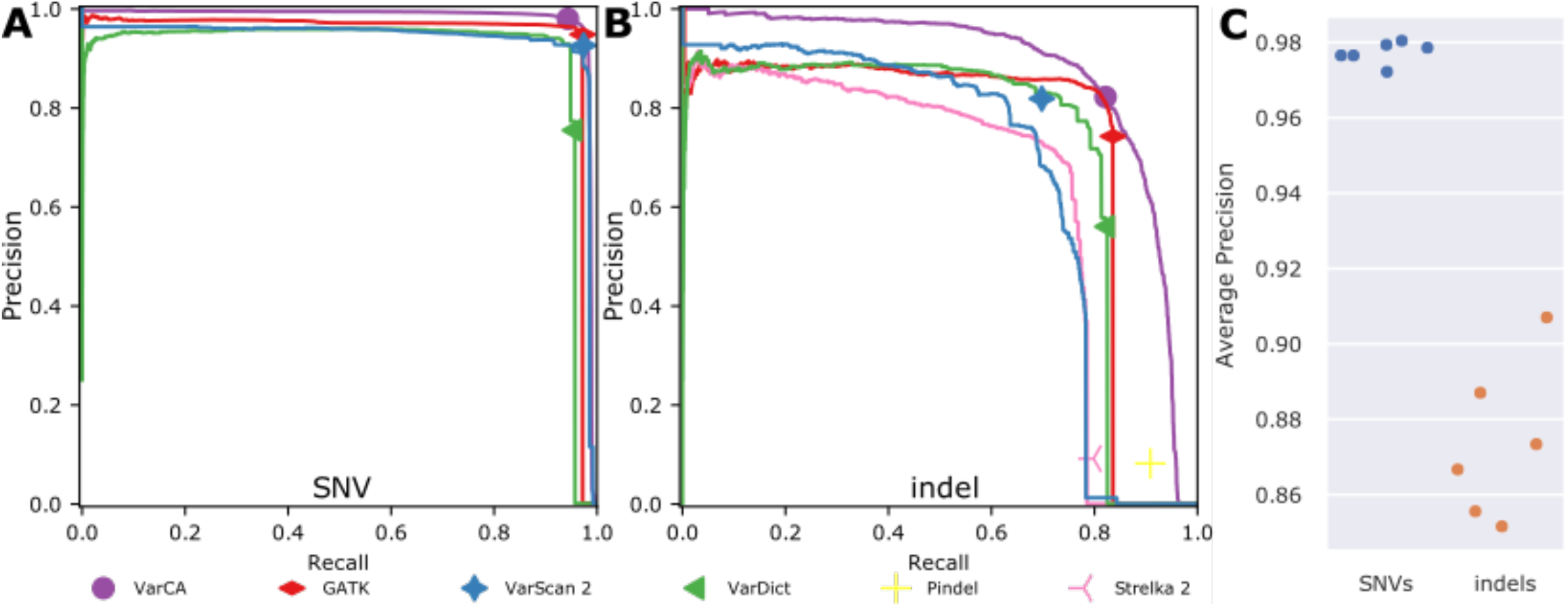
Performance of methods for detecting single nucleotide variants (SNVs) and indels on single-cell ATAC-seq. (**A**) Precision recall (PR) curves for SNV detection on reads obtained from cells that are part of single cell “cluster 12”. Overall precision and recall values for each caller, using all variant calls reported significant, are also indicated as single points. Callers that do not output a score that can be used for ranking only have a single point. The point for VarCA corresponds to a score of 0.50, which we use as a significance threshold. (**B**) PR curves for indel detection on cluster 12. (**C**) Average precision for VarCA on the 6 different clusters of human cells. VarCA is our ensemble prediction method that combines information from individual variant callers.

### VarCA feature importance

The RF classifier can evaluate how useful each feature is for predicting variants, using a metric known as importance. The importance of a feature is the decrease in the Gini impurity resulting from the use of that feature at the nodes in the trees where it is applied. We used importance to evaluate which features from each caller are most useful for SNV or indel prediction (Figs 4A and 4B and Supplementary Tables S2 and S3). For calling either type of variant, the most important feature was GATK’s QD, followed by GQ (Genotype Quality), however features from other callers also had high importance. To summarize the overall contribution from each caller, we computed a summed importance across each caller’s features. For SNVs, GATK had the highest summed feature importance, followed by VarDict and VarScan2 (Supplementary Fig S7). For indels, GATK again had the highest summed feature importance followed by VarDict, Strelka 2, and VarScan2 (Supplementary Fig S8).

**Figure 4.**
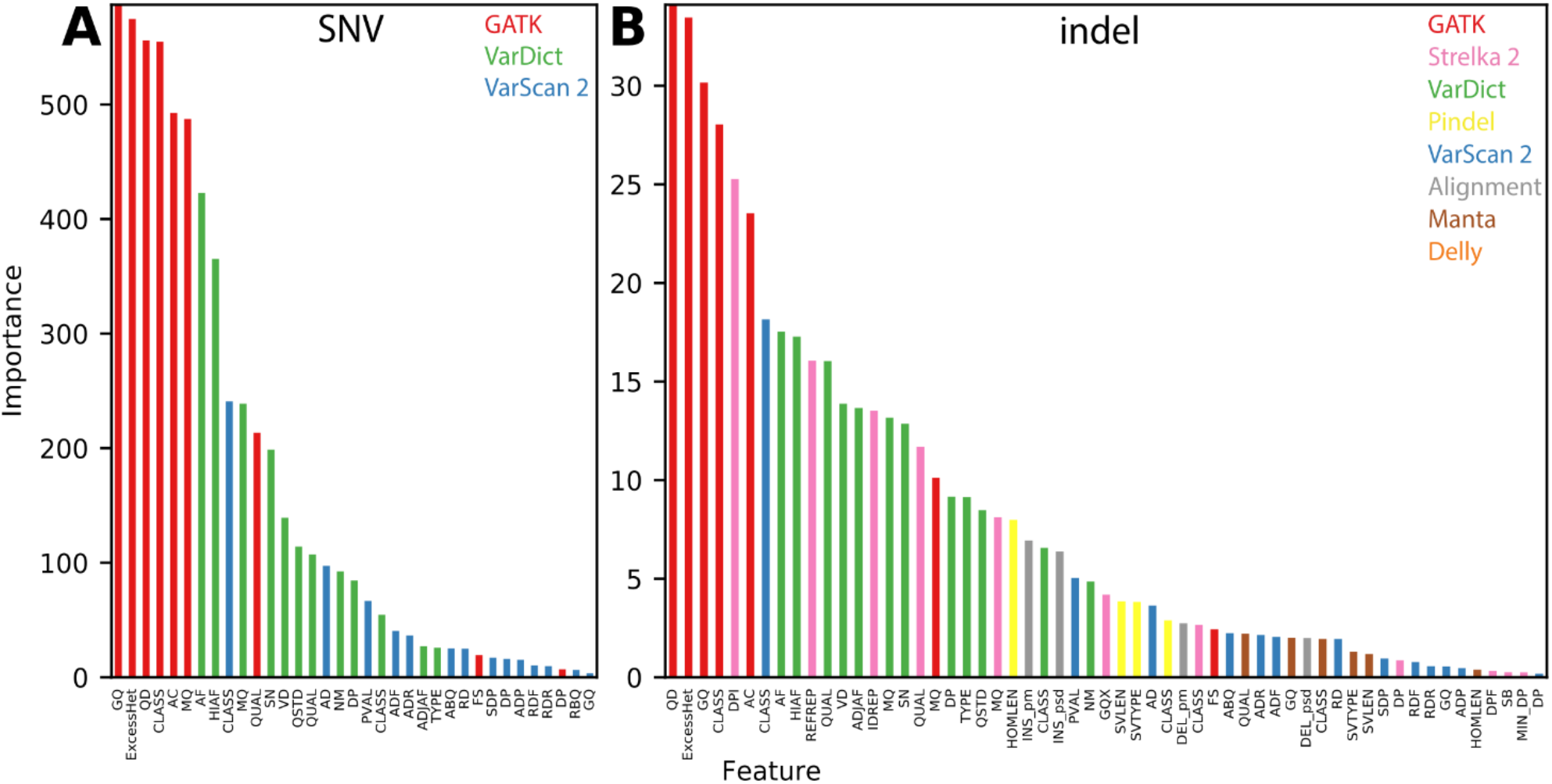
Importance of the features used by the VarCA random forest for variant classification. Colors indicate the variant caller that each feature was obtained for. These values are also provided in Supplementary Tables S2 and S3. The “Alignment” category refers to features that are extracted from read alignments by VarCA (see Supplementary Note S1) (**A**) Feature importance for single nucleotide variants. (**B**) Feature importance for insertions/deletions.

### Streamlined ensemble method

Some of the variant callers did not provide features that contributed substantially to the classification accuracy, as indicated by the summed feature importances described above (Supplementary Fig S7 and Supplementary Fig S8). We therefore considered whether these callers could be omitted from the VarCA pipeline to increase speed without sacrificing accuracy. We re-evaluated the performance of VarCA, after training on features obtained from only the top callers and found that the overall accuracy remained the same (or even improved slightly) after dropping the callers with the lowest summed feature importance (Supplementary Fig S8; Supplementary Table S9). We therefore modified the VarCA pipeline so that by default it only uses a subset of callers: GATK, VarDict, and VarScan2 for SNVs; and GATK, VarDict, Strelka 2, VarScan 2 and Pindel for indels.

### Detecting known mutations

To demonstrate the utility of VarCA for the detection of de novo mutations, we applied it to ATAC-seq data that we collected from four T-cell acute lymphoblastic leukemia cell lines (Jurkat, MOLT-4, CCRF-CEM and RPMI-8402) and one myelogenous leukemia cell line (K562). Two of these cell lines (Jurkat and MOLT-4) are known to harbor slightly different oncogenic insertions 8kb upstream of the *TAL1* gene [24] and VarCA successfully detects both of them with scores of 0.64 in Jurkat and 0.91 in MOLT-4 (Fig 5). In addition, we verify that VarCA detects an insertion associated with allele-specific expression of the *LMO2* oncogene [4] in MOLT-4 cells with a score of 0.86 (Supplementary Fig S9). The other cell lines lacked sufficient ATAC-seq coverage to call variants at these positions, except for the Jurkat and RPMI-8402 cell lines at the *LMO2* locus. For these latter cell lines, the absence of the insertion was confirmed by very low VarCA scores (Supplementary Fig S9). These results demonstrate the utility of VarCA for discovering important mutations in regulatory regions of the genome.

**Figure 5.**
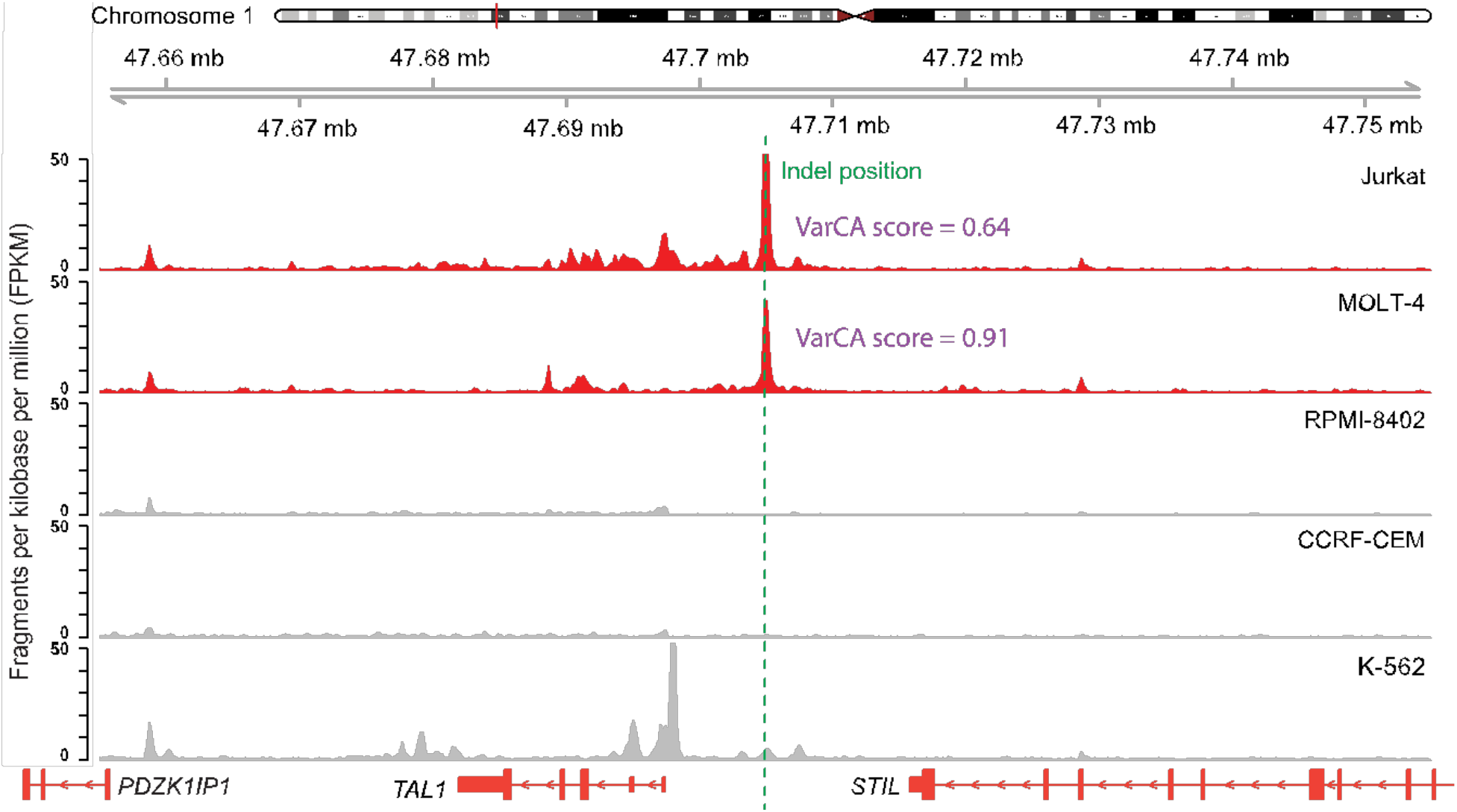
VarCA detects insertions that are known to create oncogenic enhancers upstream of *TAL1* in the Jurkat and MOLT-4 cell lines using ATAC-seq reads. The VarCA significance threshold is 0.50, and the *CCGTTTCCTAAC* insertion in the Jurkat cell line was detected with a VarCA score of 0.64. The *AC* insertion in the MOLT-4 cell line was detected with a VarCA score of 0.91.

## DISCUSSION

Using VarCA, variants can be called from ATAC-seq data in the absence of whole-genome sequencing data. By using an ensemble of variant callers, we found that SNVs and indels can be called with high accuracy within accessible chromatin regions using ATAC-seq reads. Of the individual variant callers we tested on ATAC-seq data, GATK had the highest overall performance. However, VarCA, achieves substantially better performance than any individual method and its recalibrated quality scores can be used to filter for high confidence variants.

The plots and tables presented in our results are automatically produced by the *classify* subworkflow of the pipeline. Therefore, VarCA can also be used to evaluate the performance of variant callers on ATAC-seq data in addition to those we have investigated here. Because VarCA is implemented with Snakefiles, it is easy to install, configure, and execute in a variety of Unix/Linux environments, including high-performance computing clusters.

VarCA has a few limitations. First, it expects that ATAC-seq reads will be paired-end and cannot currently be used for single-end datasets. Second, VarCA currently only supports calling of SNVs and indels, and does not detect events such as copy number alterations or inversions. Finally, a limitation of calling variants from ATAC-seq reads is that only a small fraction of the genome is covered by a sufficient number of reads. In the GM12878 ATAC-seq dataset, we called variants in ATAC-seq peaks that were covered by at least 10 sequencing reads, resulting in 4.87 MB of the genome within callable regions. While these regions of the genome are the most likely to harbor regulatory variants, mutations or variants within regulatory elements may not be detectable if they completely disrupt the regulatory region resulting in the loss of open chromatin. Nonetheless, we have found that, within the GM12878 lymphoblastoid cell line the vast majority of SNVs and indels within ATAC-seq peaks are detectable with high precision using just ATAC-seq reads.

In conclusion, we performed the first systematic evaluation of variant calling methods applied to ATAC-seq reads and found that SNVs and indels can be called with high accuracy within accessible chromatin regions for both bulk and single-cell ATAC-seq reads. Our ensemble method, VarCA, uses features output by other variant calling methods to achieve better performance than any individual method. The VarCA pipeline can also be used to evaluate the performance of any other variant callers on ATAC-seq reads. We anticipate that VarCA will be useful for the discovery of rare variants and regulatory mutations in large panels of samples that have ATAC-seq data in the absence of whole-genome sequencing [25]. Furthermore, application of VarCA to single-cell ATAC-seq datasets could potentially reveal the presence of somatic mutations that are present in only some subsets of cells.

## AVAILABILITY

The source code for VarCA is released under an MIT open source license and is available from GitHub (https://github.com/aryarm/varCA)

## ACCESSION NUMBERS

ATAC-seq data from the Jurkat, RPMI-8402, MOLT-4, CCRF-CEM and K-562 cell lines have been submitted to GEO under accession GSE129086.

## SUPPLEMENTARY DATA

Supplementary Data are available at NAR online.

## ACKNOWLEDGEMENT

We thank Nasun Hah and members of the Salk Institute Next Generation Sequencing Core for technical support with ATAC-seq experiments.

## Author’s contributions

G.M. and A.S. conceived of the project. G.M., A.R.M., A.S., and J.J. wrote the manuscript. A.S. implemented an early version of VarCA (known as BreakCA) and performed initial analyses. A.R.M. implemented VarCA and performed most of the analyses. J.J. analyzed the single-cell data with assistance from A.R.M. S.T.T. performed the ATAC-seq experiments. Y.F. performed bioinformatic processing and QC of the ATAC-seq data under the supervision of G.E. G.M. supervised the research.

