## Supplementary Materials for "Discovering single nucleotide variants and indels from bulk and single-cell ATAC-seq"

**Supplementary Material**

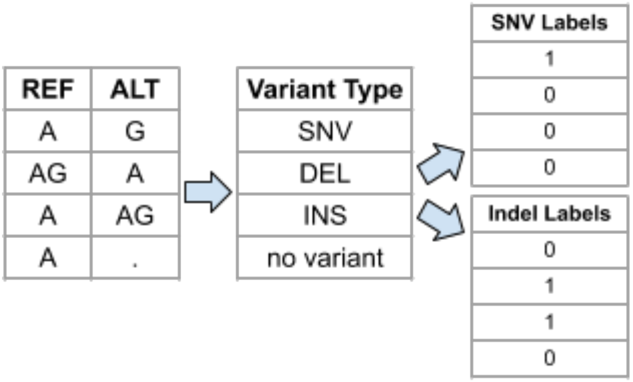


**Supplementary Figure S1.** Assigning variant labels. The random forest used by VarCA operates on binary classification labels and can predict the presence of either single nucleotide variants (SNVs) or insertions/deletions (indels). To assign labels to the truth dataset, the REF/ALT alleles are converted to binary values for both SNVs and indels. The same labeling scheme is used for the individual variant callers in the ensemble.

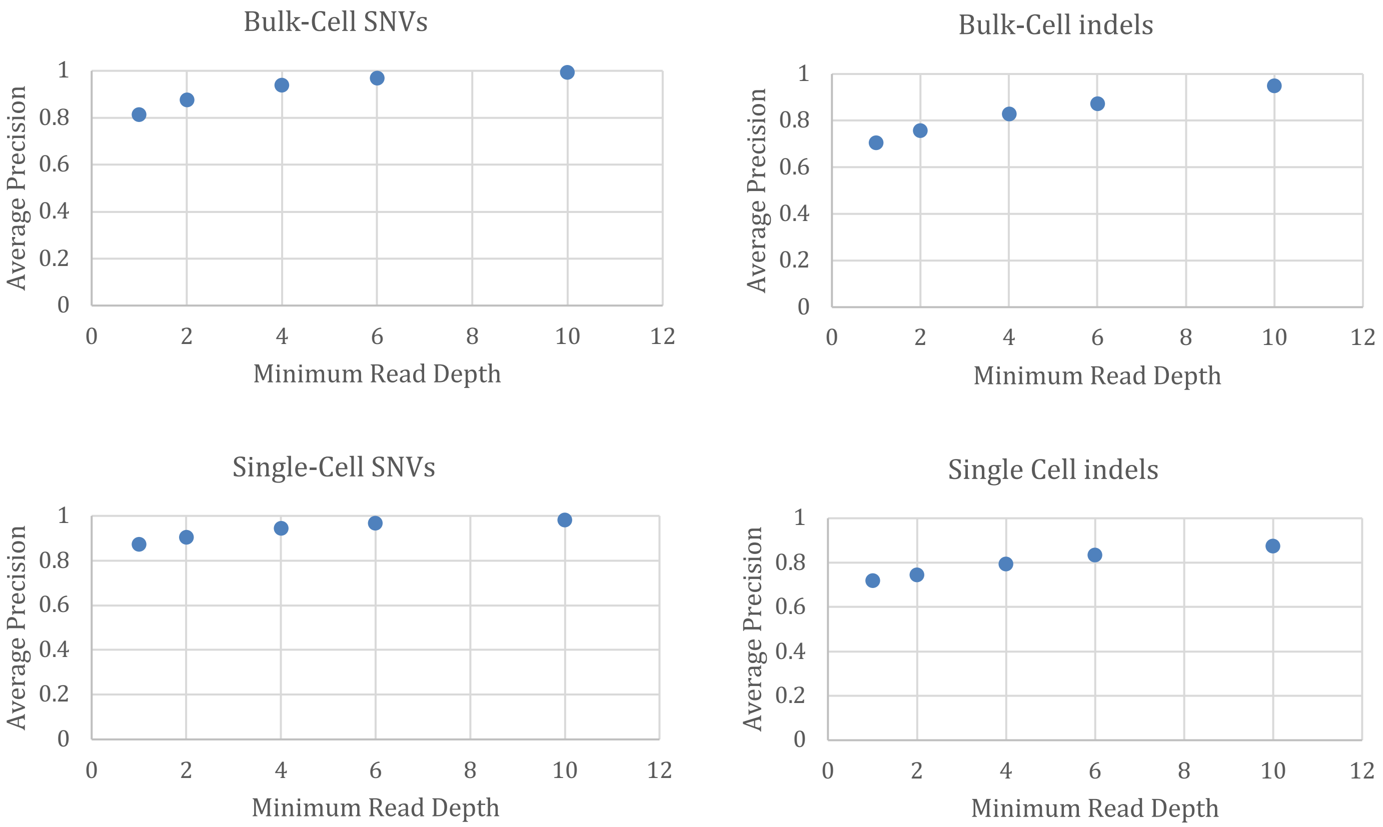


**Supplementary Figure S2.** The average precision of VarCA at different read depth thresholds when trained on bulk and single-cell ATAC-seq data for all chromosomes from the GM12878 cell line. Single-cell ATAC-seq metrics are shown for “cluster 12”.

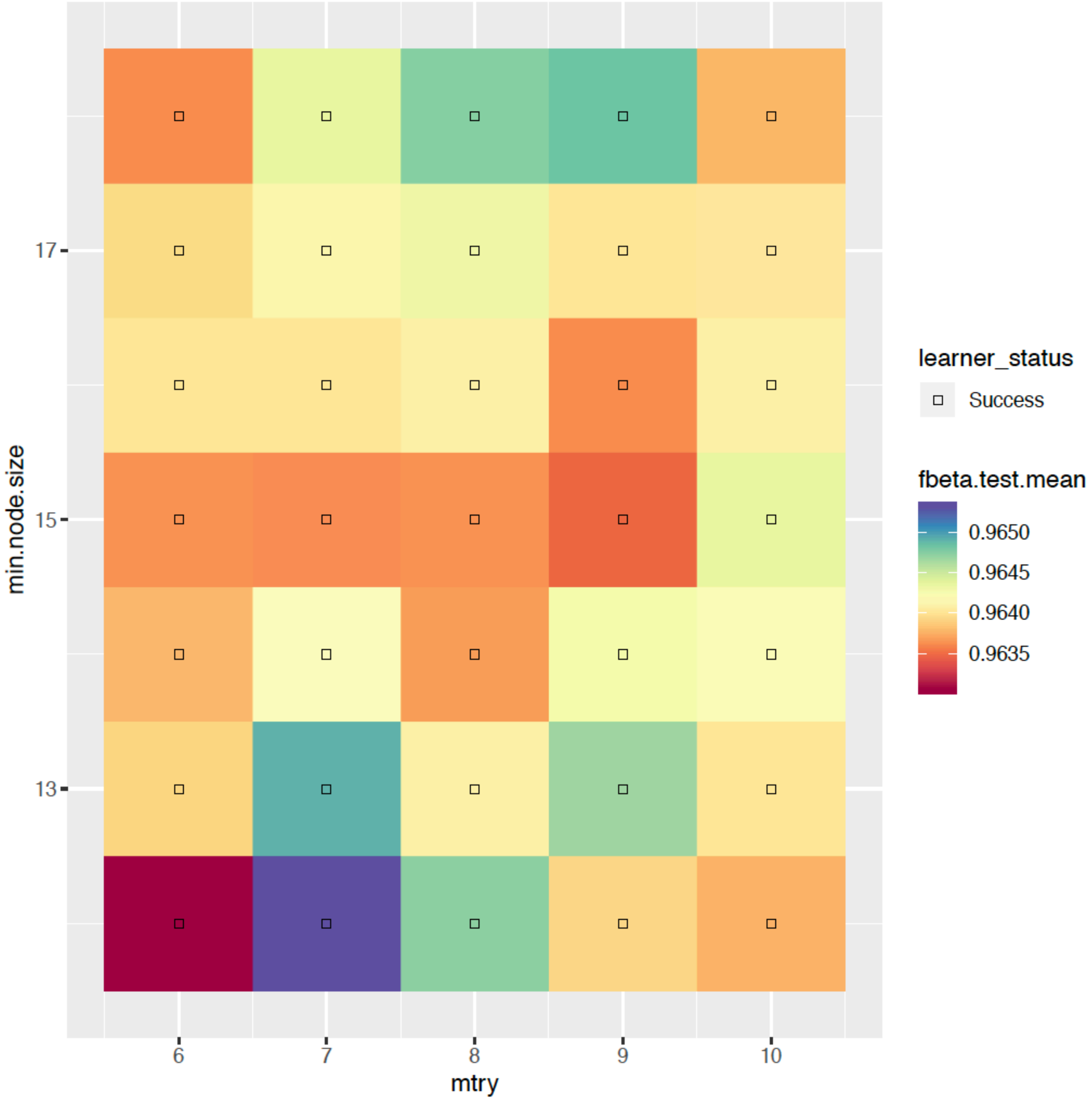


**Supplementary Figure S3.** Tuning the hyperparameters of the random forest (RF) on training data for single nucleotide variants. We trained and ran the random forest using a grid of values for the *mtry* and *min.node.size* hyperparameters that control the random forest structure. We performed 5-fold cross validation for each choice of hyperparameters and evaluated performance by computing the mean F-beta score across five cross-validation instances. We set beta to be 0.5 to weight precision higher than recall. We found that the RF performance was fairly insensitive to the choice of hyperparameter values with only small differences in F-beta.

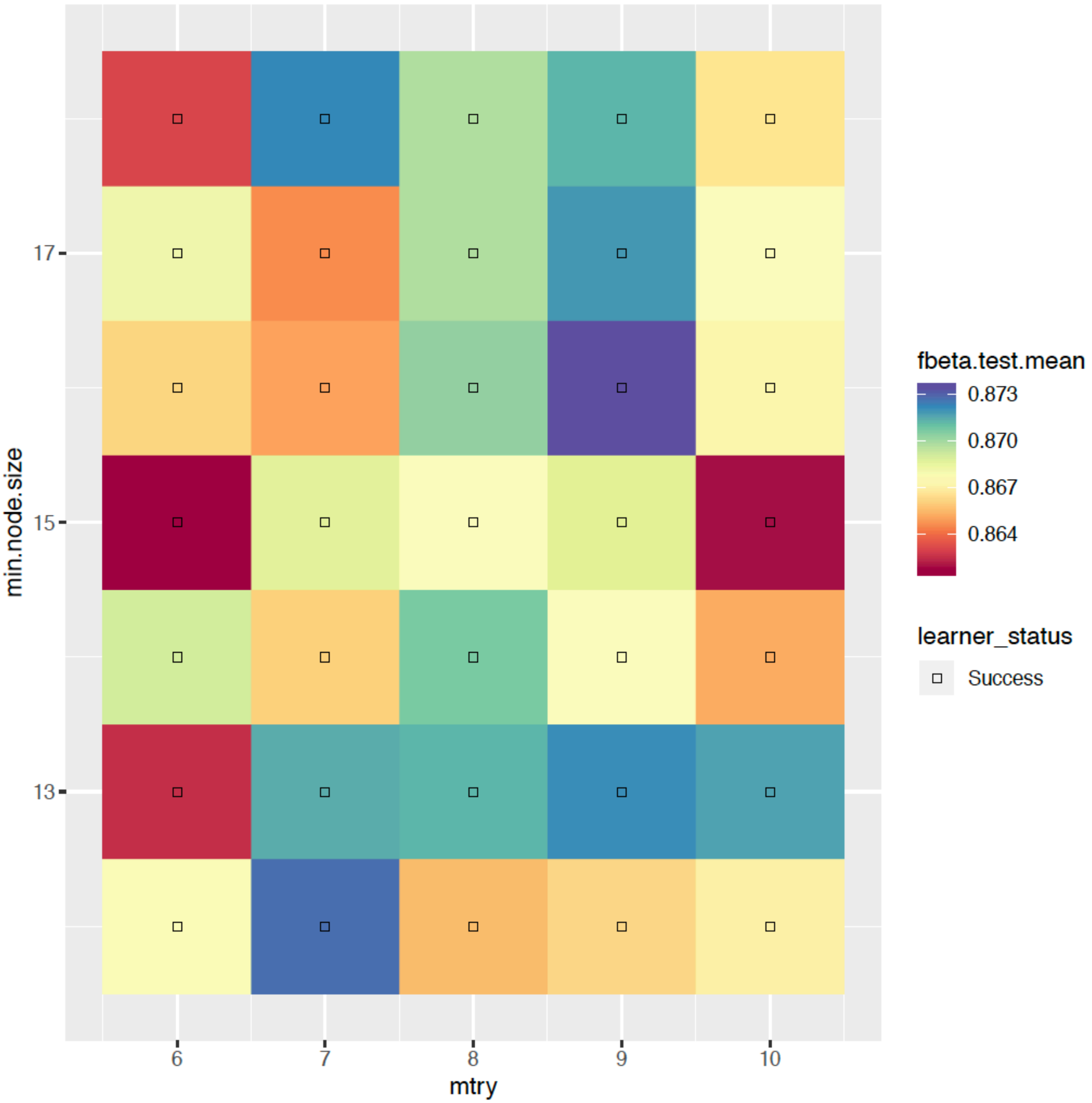


**Supplementary Figure S4.** Tuning the hyperparameters of the random forest (RF) for insertions/deletions. Performance was evaluated as described in Supplementary Figure S3.

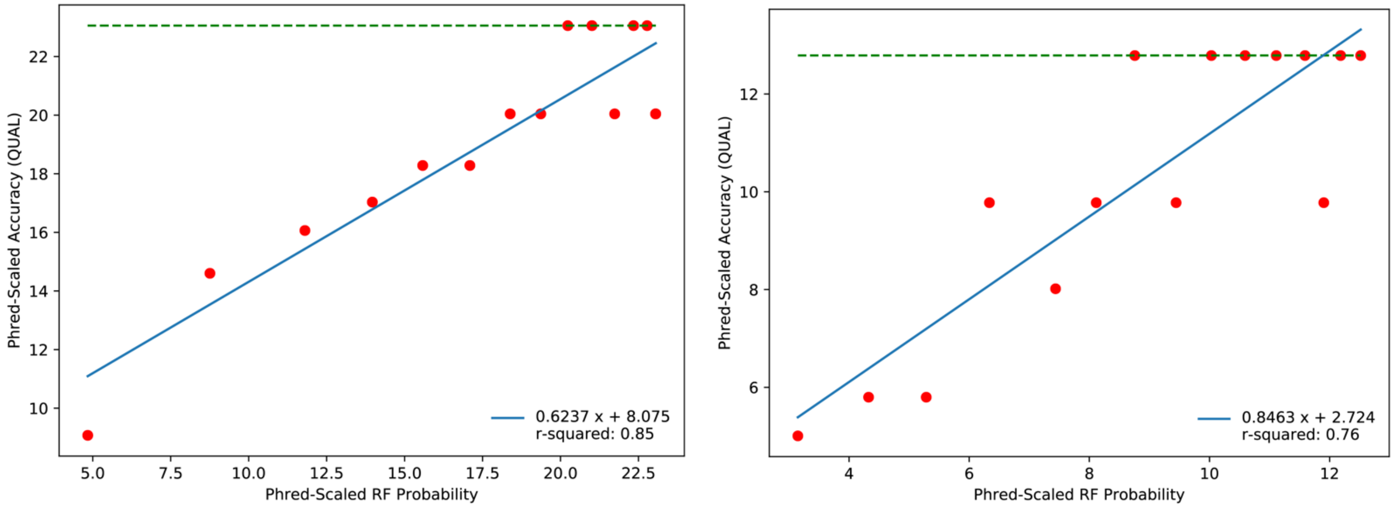


**Supplementary Figure S5.** Linear models used for converting random forest probabilities to QUAL scores for SNVs (left) and insertion/deletions (right). A QUAL score is the phred-scaled error probability of predicted variants. We computed the empirical local false discovery rate from binned predicted variants on even chromosomes of GM12878 and performed a linear regression against the mean VarCA random forest probability computed using the same bins. Predictions were performed using the VarCA random forest, which was trained on odd chromosomes. The data were ordered by the random forest probability and grouped into 15 evenly-sized bins and the empirical QUAL was calculated as $-10\log_{10}\left(1-\frac{TP}{TP+FP+c}\right)$ where $c = 1$ is a pseudo count and $TP$ and $FP$ are true and false positive counts within a bin. The maximum possible empirical QUAL score, given the number of variants in each bin, is indicated by the dashed green line.

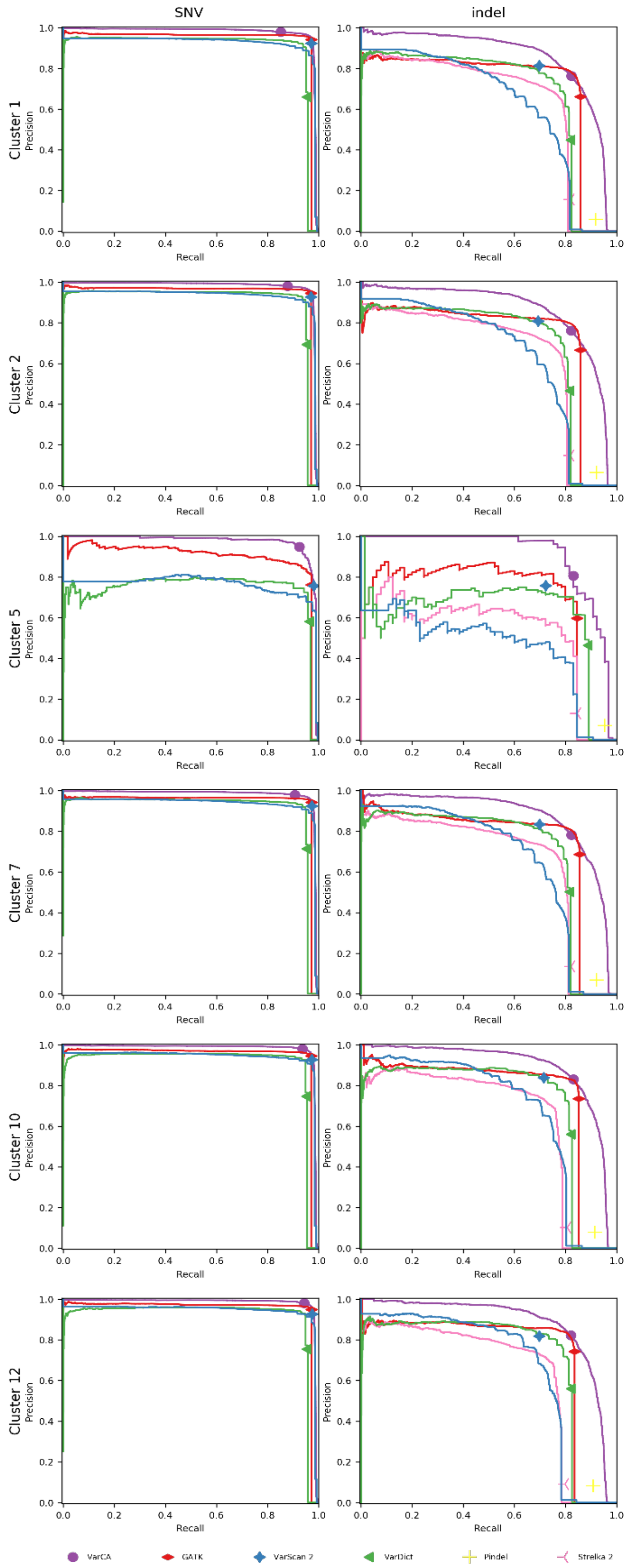


**Supplementary Figure S6.** Precision recall (PR) curves of variant callers on single-cell ATAC-seq. The callers were applied separately to the single-cell clusters corresponding to GM12878 cells (1, 2, 5, 7, 10, and 12). Only the variant callers which performed well on the bulk GM12878 were applied. The point for VarCA corresponds to a score of 0.50, which we use as a significance threshold.

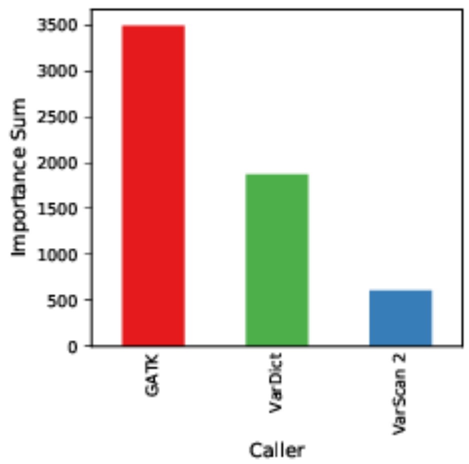


**Supplementary Figure S7.** Random forest feature importance summed by caller for single nucleotide variants.

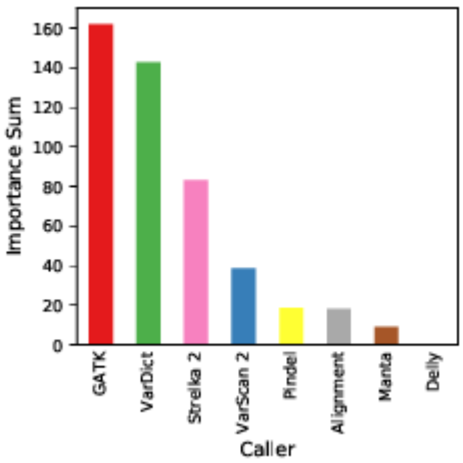


**Supplementary Figure S8.** Random forest feature importance summed by caller for insertions/deletions.

Alignment refers to features computed from the aligned reads (see Supplementary Note S1).

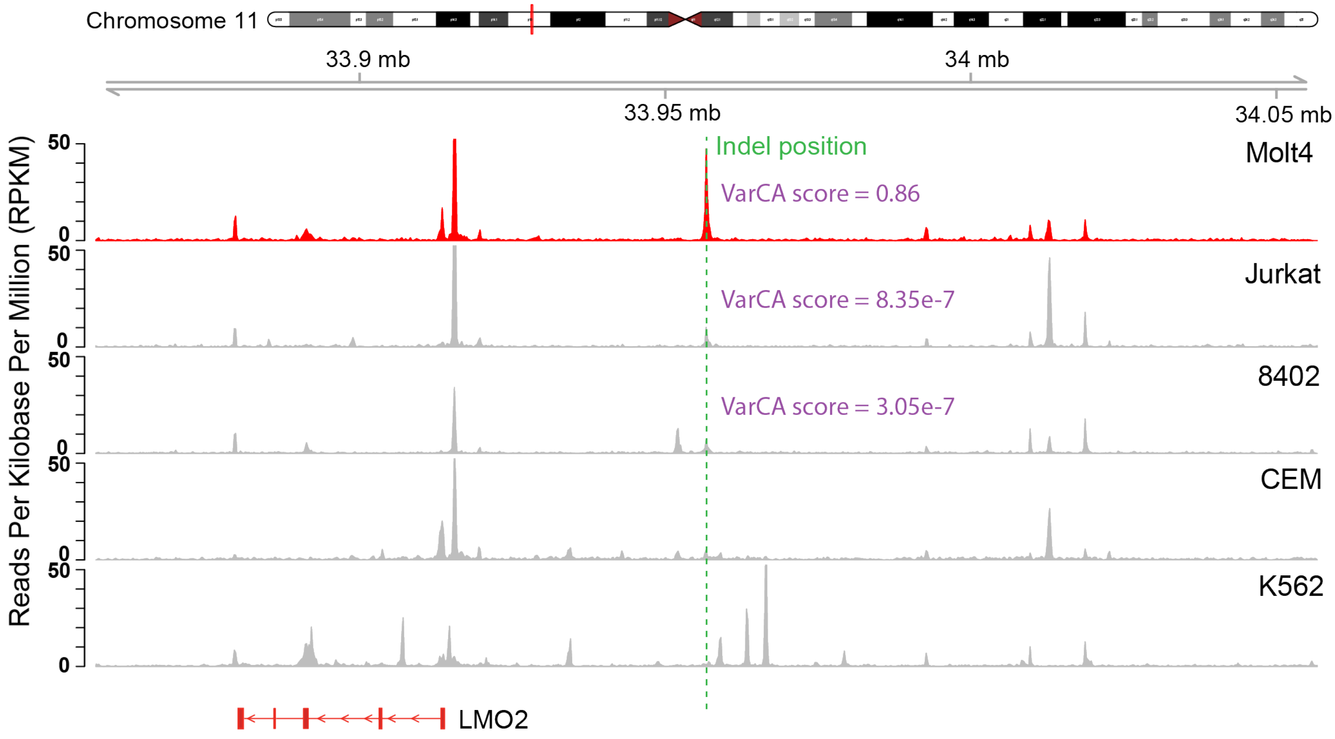


**Supplementary Figure S9.** A known oncogenic insertion upstream of *LMO2* was detected by VarCA. We applied VarCA to 5 leukemia cell lines, and the insertion was detected only in the MOLT-4 cell line, where it is known to be present, with a score of 0.86.

| caller | version | arguments | notes |
|---|---|---|---|
| GATK | 4.1.7.0 | *defaults* | HaplotypeCaller run in GVCF mode |
| VarScan | 2.4.4 | mpileup2cns --p-value 1 --strand-filter 0 | |
| VarDict-Java | 1.7.0 | -v -c 1 -S 2 -E 3 -VS SILENT | |
| DELLY | 0.8.3 | *defaults* | |
| Pindel | 0.2.5b9 | *defaults* | |
| Manta | 1.6.0 | --exome | all relevant config options were set to 0 |
| Strelka | 2.9.10 | --exome | default config options except minMapq was set to 10 |

**Supplementary Table S1.** Software versions and arguments to individual variant callers.

*(see excel file)*

**Supplementary Table S2.** Description and importance of features used by the VarCA random forest for single nucleotide variant classification.

*(see excel file)*

**Supplementary Table S3.** Description and importance of features used by the VarCA random forest for classification of insertions/deletions.

| Metric | VarCA | GATK | VarScan 2 | VarDict |
|---|---|---|---|---|
| Recall | 0.948 | 0.970 | 0.962 | 0.957 |
| Precision | 0.985 | 0.919 | 0.778 | 0.569 |
| F-Beta | 0.977 | 0.929 | 0.809 | 0.619 |
| AUROC | 0.997 | 0.985 | 0.980 | 0.978 |
| Average Precision | 0.986 | 0.925 | 0.702 | 0.705 |
| True Positives | 2983 | 3052 | 3025 | 3011 |
| False Positives | 46 | 268 | 864 | 2281 |
| True Negatives | 2311395 | 2311173 | 2310577 | 2309160 |
| False Negatives | 163 | 94 | 121 | 135 |

**Supplementary Table S4.** Metrics summarizing the performance of each single nucleotide variant caller and the ensemble method, VarCA, on bulk ATAC-seq data from the GM12878 cell line.

| Metric | VarCA | GATK | VarScan 2 | Manta | VarDict | Pindel | Strelka 2 | DELLY |
|---|---|---|---|---|---|---|---|---|
| Recall | 0.803 | 0.822 | 0.694 | 0.241 | 0.838 | 0.925 | 0.784 | 0.000 |
| Precision | 0.931 | 0.871 | 0.860 | 0.906 | 0.533 | 0.116 | 0.099 | 0.000 |
| F-Beta | 0.902 | 0.861 | 0.821 | 0.583 | 0.575 | 0.141 | 0.120 | 0.000 |
| AUROC | 0.982 | 0.911 | 0.902 | 0.617 | 0.919 | | 0.883 | |
| Average Precision | 0.896 | 0.766 | 0.386 | 0.217 | 0.471 | | 0.380 | |
| True Positives | 257 | 263 | 222 | 77 | 268 | 296 | 251 | 0 |
| False Positives | 19 | 39 | 36 | 8 | 235 | 2252 | 2291 | 144 |
| True Negatives | 2311230 | 2311210 | 2311213 | 2311241 | 2311014 | 2308997 | 2308958 | 2311105 |
| False Negatives | 63 | 57 | 98 | 243 | 52 | 24 | 69 | 320 |

**Supplementary Table S5.** Metrics summarizing the performance of each insertion/deletion caller and the ensemble method, VarCA, on bulk ATAC-seq data from the GM12878 cell line.

| Cell Line | Cell Line Type | Mapped Read Pairs (Fragments) | Fragments in peaks | FRiP | TSS Enrichment |
|---|---|---|---|---|---|
| Jurkat | T-ALL | 10,908,944 | 2,422,279 | 0.22 | 5.0 |
| RPMI-8402 | T-ALL | 27,760,815 | 5,473,477 | 0.20 | 3.7 |
| CCRF-CEM | T-ALL | 3,607,516 | 744,765 | 0.21 | 5.7 |
| K562 | CML | 27,898,577 | 3,881,592 | 0.14 | 3.4 |
| MOLT-4 | T-ALL | 10,171,311 | 1,663,657 | 0.16 | 4.1 |
| GM12878 | B Lymphoblastoid | 164,759,791 | 18,061,771 | 0.11 | 2.0 |

**Supplementary Table S6:** Mapped read and QC statistics for ATAC-seq datasets generated from the Jurkat, CCRF-CEM, RPMI-8402, MOLT-4, and K-562 cell lines. Reported mapped read fragments are de-duplicated and with chrM reads removed. The peaks used in this QC table were computed using the standard ENCODE ATAC-seq pipeline, which is more stringent than the peaks we used for variant calling (MACS2 arguments *--shift -75 --extsize 150 --nomodel --call -summits --nolambda --keep-dup all -p 0.01*). FRiP is the fraction of reads in peaks. The TSS enrichment was computed using the ATACseqQC Bioconductor package by Ou et al. 2018 using the RefSeq gene annotation. Note that FRiP and TSS enrichment values depend highly on the peak and gene annotations used. For comparison we also computed statistics for the GM12878 ATAC-seq dataset generated by Buenrostro et al. 2013.

*(see excel file)*

**Supplementary Table S7.** Metrics summarizing the performance of each SNV variant caller and the ensemble method, VarCA, when trained on all chromosomes from the bulk GM12878 cell line and tested on single cell ATAC-seq human clusters (1, 2, 5, 7, 10, and 12).

*(see excel file)*
**Supplementary Table S8.** Metrics summarizing the performance of each insertion/deletion caller and the ensemble method, VarCA, when trained on all chromosomes from the bulk GM12878 cell line and tested on single cell ATAC-seq human clusters (1, 2, 5, 7, 10, and 12).

| Excluded Callers | Recall | Precision | F-Beta | Total Positives | Total Negatives | AUROC | Average Precision |
|---|---|---|---|---|---|---|---|
| DELLY, Manta, Alignment | 0.859 | 0.982 | 0.955 | 280 | 2311289 | 1.00 | 0.965 |
| DELLY, Manta | 0.856 | 0.979 | 0.951 | 280 | 2311289 | 1.00 | 0.965 |
| DELLY | 0.856 | 0.979 | 0.951 | 280 | 2311289 | 1.00 | 0.964 |
| none | 0.859 | 0.975 | 0.95 | 282 | 2311287 | 0.998 | 0.963 |
| DELLY, Manta, Alignment, Pindel | 0.844 | 0.978 | 0.948 | 276 | 2311293 | 1.00 | 0.944 |
| DELLY, Manta, Pindel | 0.841 | 0.978 | 0.947 | 275 | 2311294 | 1.00 | 0.944 |

**Supplementary Table S9.** Metrics summarizing the performance of VarCA for finding indels after excluding certain callers. In this case, VarCA was trained on all chromosomes from GM12878. Models are ordered by F-beta, with beta set to 0.5. Compared with the full model (grey), a model excluding features from DELLY, Manta, and Alignments did not have a substantially different F-beta score and is used as the streamlined VarCA model.

**Supplementary Note S1. Read alignment features included in VarCA random forest.** In addition to features output by individual variant callers, we compute features from the read alignments. Specifically, we compute Bayesian estimates of the proportion of alignments at a position containing insertions and deletions. We use empirical Bayesian estimates to prevent over-estimation of proportions when the number of aligned reads at a position is small (i.e. this is a Bayesian alternative to using a pseudo-count). We assume that the count of reads with a given characteristic (e.g. insertion) at genomic position *i*, is a binomially-distributed random variable, $X_i$ with proportion parameter $p_i$:

$$X_i \sim \text{Binom}(n_i, p_i)$$

where, $n_i$ is the total number of reads overlapping genomic position *i*. We place a Beta prior on $p_i$:

$$p_i \sim \text{Beta}(\alpha, \beta)$$

with $\alpha$ and $\beta$ hyperparameters that describe the shape of the distribution. We estimate $\alpha$ and $\beta$ empirically using the estimated proportion mean ($\hat{\mu}$) and variance ($\hat{\sigma}^2$) computed across all positions within peaks:

$$\hat{\mu} = \frac{1}{N}\sum_{i=1}^{N}\frac{x_i}{n_i}$$

$$\hat{\sigma}^2 = \frac{1}{N}\sum_{i=1}^{N}\left(\frac{x_i}{n_i} - \hat{\mu}\right)^2$$

where $N$ is the total number of positions, $n_i$ is the number of reads overlapping position *i*, and $x_i$ is the number of reads with a characteristic at that position (e.g. insertion). The $\alpha$ and $\beta$ hyperparameters are then calculated as:

$$\alpha = \left(\frac{1-\hat{\mu}}{\hat{\delta}^2} - \frac{1}{\hat{\mu}}\right)\hat{\mu}^2$$

$$\beta = \alpha\left(\frac{1}{\hat{\mu}} - 1\right)$$

The Beta prior distribution is conjugate with the Binomial likelihood and the corresponding posterior distribution for the proportion is:

$$p_i \sim \text{Beta}(x_i + \alpha, n_i - x_i + \beta)$$

Using this distribution, we calculate the posterior mean and posterior standard deviation at each site, *i,* for the proportion of insertion and deletion alignments. We only perform posterior calculations for base positions covered by ≥ 10 reads.
